# *In Vitro* Assessment of Efficacy and Cytotoxicity of *Prunus africana* Extracts on Prostate Cancer C4-2 Cells

**DOI:** 10.1101/2021.03.14.435338

**Authors:** Peace C. Asuzu, Alberta N.A. Aryee, Nicholas Trompeter, Yasmin Mann, Samuel A. Besong, Randall L. Duncan, Carlton R. Cooper

## Abstract

Phenolic compounds are products of secondary plant metabolism known for their biological activity including their antimicrobial, antioxidant, analgesic, stimulant, anti-carcinogenic, and aphrodisiac properties. The main objective of this study was to assess the potency/cytotoxic effects of *Prunus africana* extracts on prostate cancer cells *in vitro*. Using different concentrations of *P. africana* extracts, prostate cancer C4-2 cells, a hormonally insensitive subline of LNCaP cells, were treated in a proliferation assay. A concentration dependent inhibition of cell growth in cells treated with *P. africana* bark and root extracts was present from days 1 through 3 of incubation, with the methanol extract of the bark showing the strongest effect. Compared to other plant parts, leaf extracts were significantly less cytotoxic at the same concentrations. As C4-2 cells are hormonally insensitive and designed to mimic advanced prostate cancer, crude extracts of *P. africana* are a possible treatment option, not only for hormone sensitive prostate cancer, but also advanced, hormonally insensitive prostate cancer.

## Introduction

Prostate cancer is a significant cause of morbidity and mortality worldwide and with an estimated 174,650 new cases and 31,620 deaths in 2019 according to the American Cancer Society, it is the most frequently diagnosed cancer and second most frequent cause of cancer deaths in US males (Scher, Solo, Valant, Todd, & Mehra, 2015; Shenouda et al., 2007). Approximately, 9 - 11% of men are at risk of clinically suffering from prostate cancer in their life-time globally (Komakech, Kang, Lee, & Omujal, 2017). Prostate cancer is a dynamic disease that changes over time as a function of the intrinsic properties of the tumor, patient factors, and the specific therapies to which the tumor has been exposed (Scher et al., 2015). The etiology of prostate cancer is still undecided, but a strong correlation exists with age and individuals with a family history of disease (Cunningham & You, 2015). The most consistent correlation for prostate cancer prevention is a significant consumption of fruits, vegetables, and whole grains, which are potential sources of phytoestrogens (Shenouda et al., 2007). Phytoestrogens are also found in many plants, which are commonly used in traditional medicine (Azadbakht, 2007; Joshi et al., 2011; Shenouda et al., 2007).

For years, prostate cancer has been managed through conventional treatment modalities such as surgery, radiation therapy, cryosurgery, and hormone therapies, but there is still no effective treatment for advanced stages of prostate cancer, which is hormonally resistant and bone metastatic. Chemoprevention and chemotherapy have been identified as effective approaches by which the prevalence of prostate cancer can be reduced, suppressed, or reversed (Komakech et al., 2017). Although the treatment of localized disease has significantly improved, once prostate cancer progresses to the periprostatic space by penetration and perforation of the prostate capsule and/or by invasion of the perineural spaces to the lymph nodes, few therapeutic options with no durability are available (Thalmann et al., 2000). Androgen ablation therapy, either by chemical or by surgical castration, is the last line of defense and has been gold standard for the treatment of advanced prostate cancer since Charles Huggins first pioneered this approach in 1941 (Chen et al., 2006). Although the initial response is a dramatic reduction and palliation of symptoms, prostate cancer eventually progresses to a lethal, hormone-refractory stage, for which no curative therapies currently exist (Chen et al., 2006; Thalmann et al., 2000). It has become clear that the progression from the androgen sensitive (AS) stage to the androgen-independent (AI) or hormone-refractory stage is the critical step that determines whether an individual’s disease can be cured (Chen et al., 2006).

### The History of *Prunus africana* treatment and antioxidant potential

Free radicals and oxidants generate a phenomenon called oxidative stress, a process that can change the cell membranes and other structures such as proteins, lipids, lipoproteins, and deoxyribonucleic acids (DNA) (Chandra et al., 2012; Hajhashemi, Vaseghi, Pourfarzam, & Abdollahi, 2010; Liu et al., 2018; Pham-Huy, He, & Pham-Huy, 2008). In humans, oxidative damage and free radicals are associated with a number of chronic diseases including cancer (Hajhashemi et al., 2010; Liu et al., 2018; Pham-Huy et al., 2008). In the last decades, several plants have been confirmed to contain chemo-preventive and therapeutic agents for various cancers including prostate cancer (Komakech et al., 2017; Shenouda et al., 2007). Among plants with anti-prostate cancer potential, researchers identified *Prunus africana* (African cherry), with its unique combination of phytochemicals and *Momordica charantia* (Komakech et al., 2017; Shenouda et al., 2007).

*P. africana* has been used in African traditional medicine to treat prostate cancer and related conditions across various communities for many years (Komakech et al., 2017). Previous studies have confirmed the effectiveness of the bark extract of *P. africana* in prostate disorders and attributed this to the synergistic effects of pentacyclic triterpenoids, ferulic esters of long-chain fatty alcohols, and phytosterols (Karan, Kumar Jena, Sharma, Vasisht, & Efferth, 2017; Komakech et al., 2017; Nyamai et al., 2015). Researchers have discovered that polyphenols are very good antioxidants capable of neutralizing the destructive reactivity of undesired reactive oxygen/nitrogen species produced as byproducts during metabolic processes in the body (Pandey & Rizvi, 2009). Also, epidemiological studies have revealed that polyphenols provide a significant protection against development of several chronic conditions such as cardiovascular diseases (CVDs), cancer, diabetes, infections, aging and asthma (Pandey & Rizvi, 2009). We hypothesize, therefore, that phenolic compounds in selected medicinal plants play a significant role in chronic disease prevention and chemotherapy and set out to test this hypothesis *in vitro*, with the goal of finding new drugs for prostate cancer.

Various bioactive substances with anti-inflammatory, anti-cancer, and anti-viral properties have been identified in different members of the genus *Prunus* (Jeruto, Mutai, Catherine, & Ouma, 2011; Kadu et al., 2012; Ngule, Ndiku, & Ramesh, 2014; Schleich, Papaioannou, Baniahmad, & Matusch, 2006b). *P. africana* has great medicinal value, as it is used traditionally in the treatment of a wide range of clinical conditions such as chest pain, fever, malaria, stomach ache, diarrhea, allergies, kidney disease, epilepsy, arthritis, hemorrhage, hypertension, sexually transmitted diseases, and benign prostatic hyperplasia (BPH) (Das, 2017; Jeruto et al., 2011; Komakech et al., 2017). *P. africana* extracts have also been reported to have anti-bacterial and anti-fungal activity, and the bark is used for liver problems and constipation (Jeruto et al., 2011; Madivoli et al., 2018; Ngule et al., 2014). However, the tree is predominantly used by brewing bark extracts to treat BPH, a disorder of the prostate, common in older men. The extracts significantly improve urologic symptoms having anti-proliferate and apoptotic effects on the prostate (Kadu et al., 2012; Komakech et al., 2017). *P. africana* is one of the many medicinal plants containing large quantities of bioactive compounds that can be used for prostate cancer management. Its use in cancer chemotherapy and chemoprevention has been discussed in a few peer reviewed journal articles (Boulbès et al., 2006; Komakech et al., 2017; Roell & Baniahmad, 2011; Schleich, Papaioannou, Baniahmad, & Matusch, 2006a; Shenouda et al., 2007).

### Phytochemical constituents of *Prunus africana*

The pharmacological efficacy of *P. africana* is believed to be due to various known and unknown compounds (Kadu et al., 2012; Komakech et al., 2017; Nyamai et al., 2015; Shenouda et al., 2007). Among the known compounds, three groups are of great importance: (1) phytosterols, especially β-sitosterol, with its anti-inflammatory properties that inhibit the swelling of the prostate gland (Kadu et al., 2012; Komakech et al., 2017; Nyamai et al., 2015); (2) pentacyclic triterpenoids that are anti-edematous by inhibiting glucosyl transferase activity (Kadu et al., 2012; Nyamai et al., 2015); and (3) ferulic acid esters, or their chemical derivatives, which inhibit angiogenic pathways, thus preventing the growth of new blood vessels from preexisting vessels, as well as the growth and spread of prostate cancer (Komakech et al., 2017; Nyamai et al., 2015).

Other secondary metabolites from this species include β-sitostenone, campesterol, n- tetracosanol and n-docosanol, myristic acid, linoleic acid, lauric acid, methyl myristate, methyl laurate, methyl linoleate, lup-20(29)-en-3-one, palmitic acid, (3.β.,5.α)- stigmast-7-en-3-ol, stigmastan-3,5-diene, α-tocopherol, cyanidin-O-galactoside, cyanidin-3-O-rutinoside, procyanidin B5, and robinetinidol-(4-α-8) catechin-(6,4-α) robinetinol (Mugaka et al., 2013; Nyamai et al., 2015). Phytochemicals from *P. africana*, suggested for the treatment of prostate cancer, include ursolic acid, oleanolic acid, *β*-amyrin, atraric acid (AA), N-butylbenzene- sulfonamide (NBBS), *β*-sitosterol, *β*-sitosterol-3-O-glucoside, ferulic acid, and lauric acid. Previous studies suggested that *P. africana* extracts target fast dividing prostate cancer cells by impairing mitosis or causing target cells apoptosis (Komakech et al., 2017).

*P. africana*’s anti-prostate cancer phytochemicals can be divided into three major categories based on their targets and pharmacological effects: (i) phytochemicals that kill the tumor cells through apoptotic pathways, a common mode of action of chemotherapeutic agents against a wide variety of cancer cells, (ii) phytochemicals that alter the signaling pathways required for the maintenance of prostate cancer cells, and (iii) phytochemicals that exhibit strong antiandrogenic and antiangiogenic activities (Komakech et al., 2017).

An AI cell line, C4-2, reproducibly and consistently follows the metastatic patterns of hormone-refractory prostate cancer by producing lymph node and bone metastases when injected either subcutaneously or orthotopically in either hormonally intact or castrated hosts. This model permits the study of factors that determine the tropism of prostate cancer cells for the skeletal microenvironment (Thalmann et al., 2000).

The main aim of this study was to determine the cytotoxicity of *Prunus africana* extracts on prostate cancer C4-2 cells.

## Materials and Methods

### Materials

Bark, root and leaf plant parts of *Prunus africana* were obtained from Cameroon and supplied by Dr. Besong. Dr. Robert A. Sikes of the University of Delaware donated the C4-2 prostate cancer cell lines. Roswell Park Memorial Institute (RPMI) media, trypsin/EDTA, phosphate buffered saline (PBS), dimethyl sulfoxide (DMSO) and ethanol were purchased from Sigma-Aldrich (St. Louis, MO) and Thermo Fisher Scientific (Waltham, MA) and ethanol was purchased from Decon Labs (King of Prussia, PA).

## Methods

### Sample preparation and extraction

After the plant parts from the selected medicinal plants were pulverized in a coffee grinder the particles that could pass through a screen with apertures of 1.3 mm were labelled and stored at 4°C until needed. A 10 g sample of each plant part was homogenized at a ratio of 1:10 (w/v) with one of five solvents: acetone (ACE), dichloromethane (DCM), methanol (MET), ethanol (ETH) or water (H_2_O). The slurry, stirred at room temperature on an Orbi Shaker (Benchmark, Edmonton, AB) for 24 h at 140 rpm, was subsequently filtered using a cheesecloth. The filtrate was then centrifuged at 1200 rpm for 20 min before the supernatant was decanted carefully into pre-weighed aluminum drying pans and evaporated to dryness under a fume hood. The dried extracts were weighed and stored at 4°C until needed.

### Determination of Total Phenolic Content

The total phenolic content was determined as per the method of Taga, et al. (1984). In brief, 5 mg of each extract was dispersed in 1 mL of 60:40 acidified methanol: water buffer solution, from which 100 µL was drawn, mixed thoroughly with 2 mL of 2% sodium carbonate solution (Na_2_CO_3_) and incubated at room temperature for 2 min. To this mixture (and a blank containing 100 µL of buffer solution mixed with 2 mL of Na_2_CO_3_) was then added 100 µL of 50% folin-ciocalteau reagent, and further incubation at room temperature for 30 min. The absorbance of the samples and different concentrations of gallic acid standard in the range of 0.05-0.11 mg/mL were measured at 750 nm using a DU 720 General-purpose Spectrophotometer (Beckman Coulter Inc., Brea, CA) and the concentration of phenolic compounds in the extracts was calculated from the Gallic acid standard curve and expressed as milligrams of Gallic acid equivalent per gram of extract (mg GAE/g).

### Determination of Total Flavonoid Content

Total flavonoid content was determined as per the method of Stankovic, M.S, 2011. In brief, to 4 mg of each extract dispersed in 1 mL of methanol was added 1 mL of 2% aluminum chloride solution, before incubation at room temperature for 1 h. The absorbance of the samples and the rutin standard of concentration range 0.005-0.03 mg/mL were measured at 415 nm using a DU 720 General-purpose Spectrophotometer (Beckman Coulter Inc., Brea, CA). The flavonoid content was calculated from the rutin standard curve and expressed as mg Rutin per gram of extract.

### Determination of EC_50_ and ARP

The concentration required to obtain a 50% antioxidant effect or 50% effective concentration (EC_50_) was calculated using the % DPPH remaining method. Ascorbic acid standard in six different concentrations ranging from 0.05 – 0.3 mg/mL was mixed with 0.12 mg/mL of DPPH in methanol solution and the mixture incubated in the dark, at room temperature for 30 min. The absorbance was measured under dim lighting at 515 nm using a DU 720 General-purpose Spectrophotometer (Beckman Coulter Inc., Brea, CA) and a standard curve was plotted. The EC_50_ of the plant samples were calculated using the quadratic equation of the ascorbic acid standard curve and the ARP is the reciprocal of the EC_50_. Lower EC_50_ values indicate a stronger antioxidant effect than higher values.

### Cell culture and Growth Inhibition assay

The C4-2 cells were grown to confluency in a T-75 tissue culture flask in Roswell Park Memorial Institute (RPMI) 1640 medium (Corning) supplemented with glutamine, 10% fetal bovine serum (Atlas Biologics) and antibiotics (100 µg/mL penicillin, 100 μg/mL streptomycin, Hyclone). Cells at a seeding cell density of 8 × 10^5^ cells per well were plated into six well plates containing 2 mL of medium, with sufficient plates for a 4-day proliferation study. The cells were then incubated overnight at 37°C in 5% CO_2_. The *Prunus* extracts were first dissolved in 100% ethanol or 1% dimethyl sulfoxide and further diluted with RPMI 1640 medium for the cytotoxicity assay.

Different concentrations of the extracts were tested for effect in a preliminary assay with concentrations ranging from 1 mg/mL to 0.001 mg/mL. Based on the results of the preliminary testing, extracts at concentrations of 0.025 mg/mL, 0.01 mg/mL, 0.005 mg/mL, 0.0025 mg/mL and 0.001 mg/mL of methanol and ethanol bark and root extracts (strongest antioxidant effect) were then added in triplicate to the cells and incubated for 24 h at 37°C with an ethanol/ DMSO vehicle control. After 24 h, media from one set of plates was removed, and the cells were stripped and counted using a NovoCyte flow cytometer (ACEA Biosciences, San Diego, CA), while the media from the remaining set of cells were removed and replaced with the same concentration of extracts in 2 mL of media. This was repeated every 24 h up to the 4^th^ day and a proliferation profile of the treated cells was plotted and compared to the profile of the control cells. The experiment was repeated three times and statistical significances were determined by p values ≤ 0.05.

## Results and Discussion

Bark extracts of *P. africana* showed the highest total phenolic content (TPC), while the leaf extracts showed the highest total flavonoid content (TFC), although most plant part extracts tested yielded significant TPC and TFC values (Table 1). Methanol was the most effective solvent for the TPC with the highest value being recorded for *P. africana* bark (1397.33 mg/mL), while dichloromethane yielded very low values across all samples tested (3.30 mg/mL for *P. africana* bark). A similar pattern was seen in the EC_50_ assay, where the strongest antioxidant effect was seen in the methanol bark extract of *P. africana*, a value even lower than that recorded for the same concentration of ascorbic acid (0.10 and 0.18 for methanol bark and ascorbic acid respectively). Based on these results, methanol extracts were chosen for the proliferation assay, to be compared with another polar solvent ethanol, to determine which of the two had a greater effect on the prostate cancer C4-2 cells.

**Table 1:** Total phenolic content and total flavonoid content of *P. africana*.

| Plant | Part | Extraction solvent | Total Phenolic Content (mg GAE/g) | Total Flavonoid Content (mg RU/g) |
| --- | --- | --- | --- | --- |
| *Prunus africana* | Bark | Acetone | 1348.00±35.55 | 25.13±0.81 |
| | | Dichloromethane | 3.30±0.38 | 1.45±0.43 |
| | | Methanol | 1397.33±36.90 | 17.84±0.58 |
| | | Ethanol | 982.66±11.55 | 38.63±1.23 |
| | | Water | 1054.67±13.32 | 7.85±0.37 |
| | Leaf | Acetone | 82.67±2.31 | 118.06±1.34 |
| | | Dichloromethane | 66.67±4.62 | 61.71±1.34 |
| | | Methanol | 349.33±2.30 | 55.00±1.34 |
| | | Ethanol | 178.66±2.30 | 76.02±4.31 |
| | Root | Acetone | 316.00±8.00 | 6.44±0.52 |
| | | Dichloromethane | 16.00±3.46 | 4.36±0.35 |
| | | Methanol | 430.66±6.11 | 8.67±0.16 |
| | | Ethanol | 584.00±10.58 | 6.79±0.27 |

Values are mean of three determinations ( $n = 3$ ) ± standard deviation (SD). Values in the same column (in the same block) followed by the same letter are not significant at $p \leq 0.05$

A concentration-dependent cytotoxicity was observed in the preliminary testing of bark and root extracts on C4-2 cells, with the methanol extract of the bark showing a stronger effect than the others. There was complete lysis of all cells above 0.05 mg/mL for all bark and root samples. Differences between the different extracts was most obvious at 0.05 mg/mL (Fig. 1A- G). A proliferation effect was also observed, especially in concentrations below 0.01 mg/mL when incubated for more than 24 h. This may be due to some previously undescribed compounds in the extracts or a different composition of the same compounds in this sample of *P. africana*, resulting from its genetic composition and epigenetic factors. Compared to other plant parts, leaf extracts showed much less cytotoxicity at the same concentrations, although there was marked clustering of cells and proliferation compared to the control. This may indicate a higher percentage of the compound or combination of compounds responsible for the proliferation seen with treatment of cells using bark and root extracts. In support of this hypothesis, Mugaka et al. (2013) reported differences in secondary metabolite (β-sitosterol and n-docosanol) content between the different plant parts of *P. africana* (bark and leaves) and differences between plants from different geographical regions in Tanzania (Mugaka et al., 2013).

**Figure 1:**
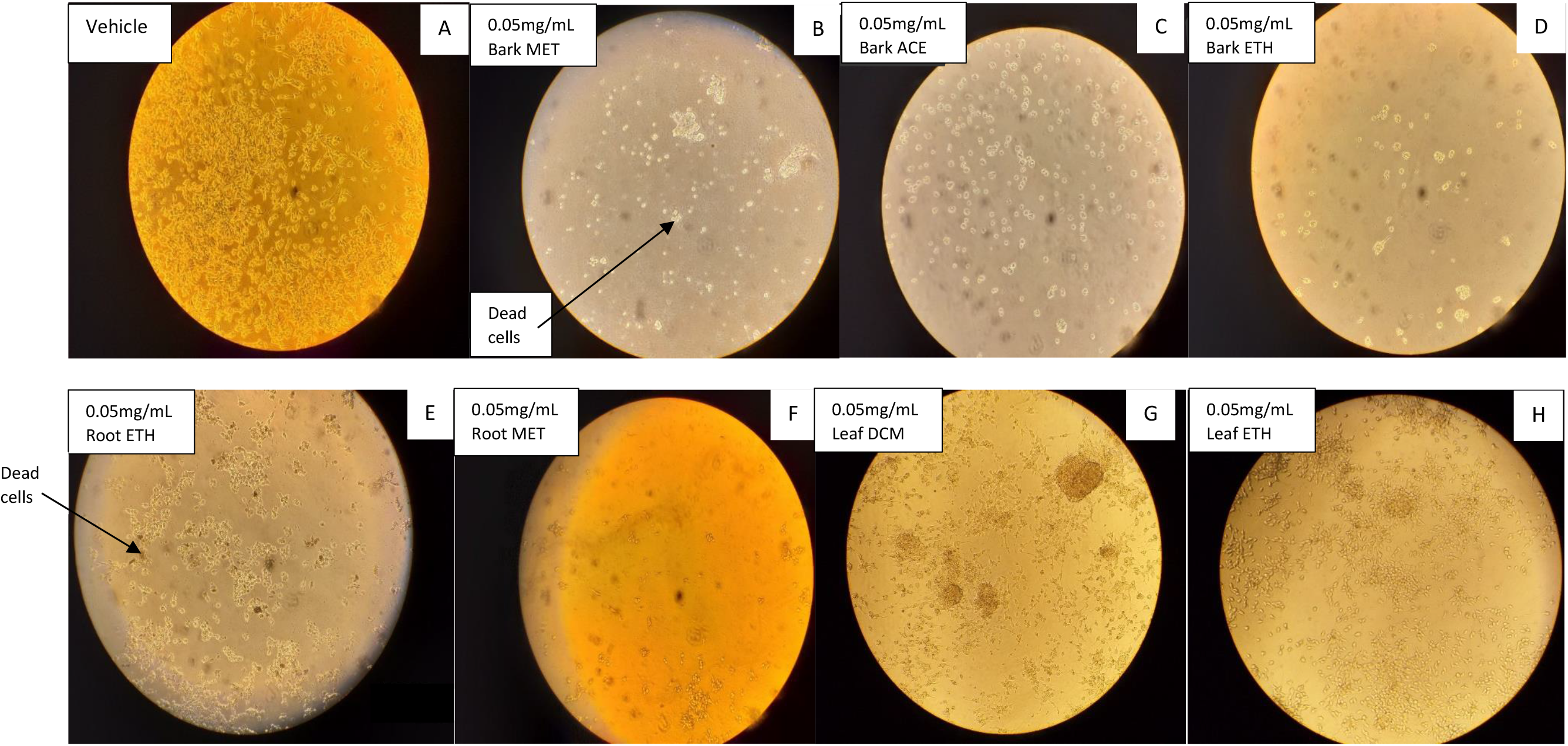
C4-2 cells after 24 h exposure to treatment with different *P. africana* plant part extracts. Cells treated with ethanol in media (control) (A), 0.05 mg/mL *Prunus* bark methanol extract (B), *Prunus* bark acetone extract (C), *Prunus* bark ethanol extract (D), *Prunus* root ethanol extract (E), *Prunus* root methanol extract (F), *Prunus* leaf dichloromethane extract (G) and *Prunus* leaf ethanol extract (H).

Mostly dead cells are visible with some granulation-like material in areas of cell clusters in Fig. 1B. Most cells in Fig. 1C are round but still appear attached to the bottom of the well. Many dead or round cells are present in Fig. 1D, with a few healthy-looking cells as well. Most cells in Fig. 1E are dead or round with significant debris but a few healthy-looking cells are present. Most cells in Fig. 1F are also dead or round with a few healthy-looking cells present, but healthy cells are fewer than those seen in Fig. 1E. Very little cell death is seen in Figs. 1G and H compared to cells treated with extracts from the bark and root. However, there is marked cell clustering, and rapid proliferation appears to be occurring in both treatments.

The phenomenon hormesis, which is a dose-response phenomenon characterized by a low-dose stimulation and a high-dose inhibition may account for the proliferation seen at lower doses (V. Calabrese et al., 2010). Southam and Ehrlich first presented the term hormesis in published literature in 1943 by, reporting that low doses of extracts from the red cider tree enhanced the proliferation of fungi with the overall shape of the dose response being biphasic. Similar types of dose-response observations have been subsequently reported by other researchers (E. J. Calabrese, 1999, 2005; V. Calabrese et al., 2010). This low-dose stimulation that manifests immediately below the pharmacological and toxicological thresholds is modest in magnitude, being at most about 30 - 60% greater than the control group response and may result from either a direct stimulation or via an overcompensation stimulatory response after disruption in homeostasis (E. J. Calabrese, 1999; V. Calabrese et al., 2010). Hormetic-like biphasic dose responses have been reported and demonstrated in 136 tumor cell lines from over 30 tissue types for over 120 different agents (E. J. Calabrese, 2005; V. Calabrese et al., 2010).

*In vitro* cytotoxicity screening is a valuable tool in drug discovery and is used widely by researchers, especially when bio-prospecting for potentially active cancer drugs (Maiyo, Moodley, & Singh, 2016). Results from the proliferation assay showed growth inhibition of C4-2 cells by *P. africana* bark and root extracts in a dose dependent manner. Shenouda et al. (2007) reported similar results in a tissue culture study, where they suggested that ethanolic extracts of *P. africana* bark inhibited the growth in human prostate cancer cell lines (PC-3 and LNCaP) by 50% at 2.5 µL/mL and induced significant apoptosis in both cell lines (PC-3 and LNCaP) at 2.5 µL/mL compared to untreated cells (Shenouda et al., 2007). This shows that crude extracts of *P. africana* inhibit growth of both hormonally sensitive (LNCaP) and hormonally insensitive (C4-2) cells modelling advanced prostate cancer at similar concentrations. Thus, the crude extract has the potential for treatment of advanced prostate cancer which is hormonally insensitive. Although there was appreciable growth inhibition in this study, there was a significant difference only between the control cells and cells treated with the highest concentration (0.025 mg/mL) of bark methanol extract (p <0.05) (Figs. 17 and 18), indicating the need of a significant dose to control advanced stage prostate cancer cells.

The inhibition of the growth of C4-2 cells was present from day 1 through day 4 of incubation with the *P. africana* extract as seen in the study by Shenouda et al. (2007) but there were variations in the percentages of inhibition. The maximum inhibition by treatment with 0.025 mg/mL methanol and ethanol bark extracts was seen on day 2 (82% and 30% reduction, respectively), and for methanol root extract on day 3 (37% reduction), compared to the control, unlike in the previous study where a more consistent inhibition was seen (50%) compared to the control. This may be due to differences in metabolite composition between bark and root extracts and methodology as the previous study assayed for the total DNA concentration using the thymidine incorporation assay, while simple cell counting was used in this study (Shenouda et al., 2007). In addition, C4-2 cells differ from the cell lines used in the previous study by being hormone insensitive. However, it is clear that the bark methanol extract showed more potent inhibition of cell growth than did the root methanol extract Figs. 2 and 4.

**Figure 2:**
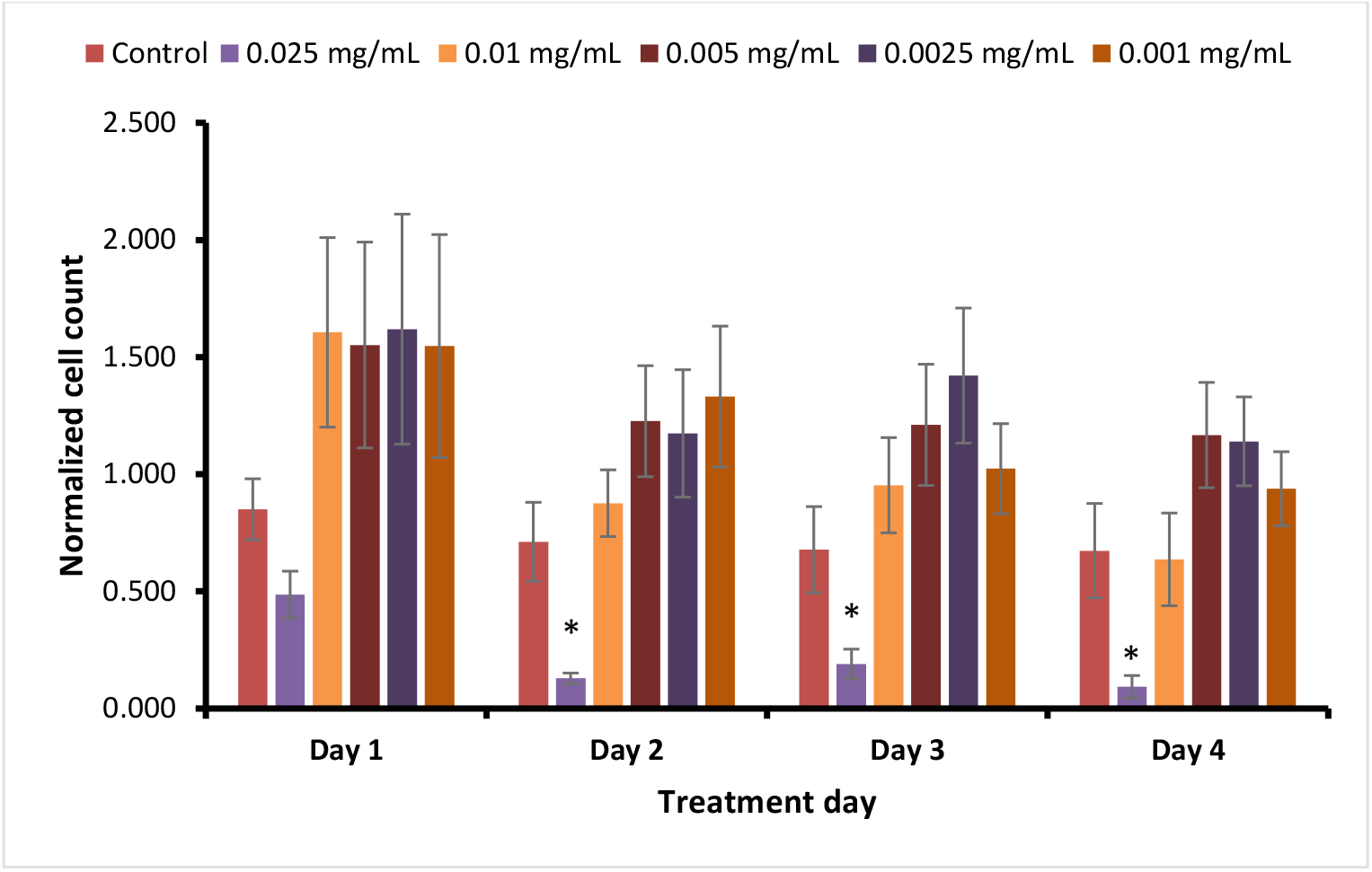
Proliferation assay for *P. africana* bark methanol extract. Values are mean of three determinations (n = 3) ± SE. Asterisk (*) denotes the presence of statistical significance.

**Figure 3:**
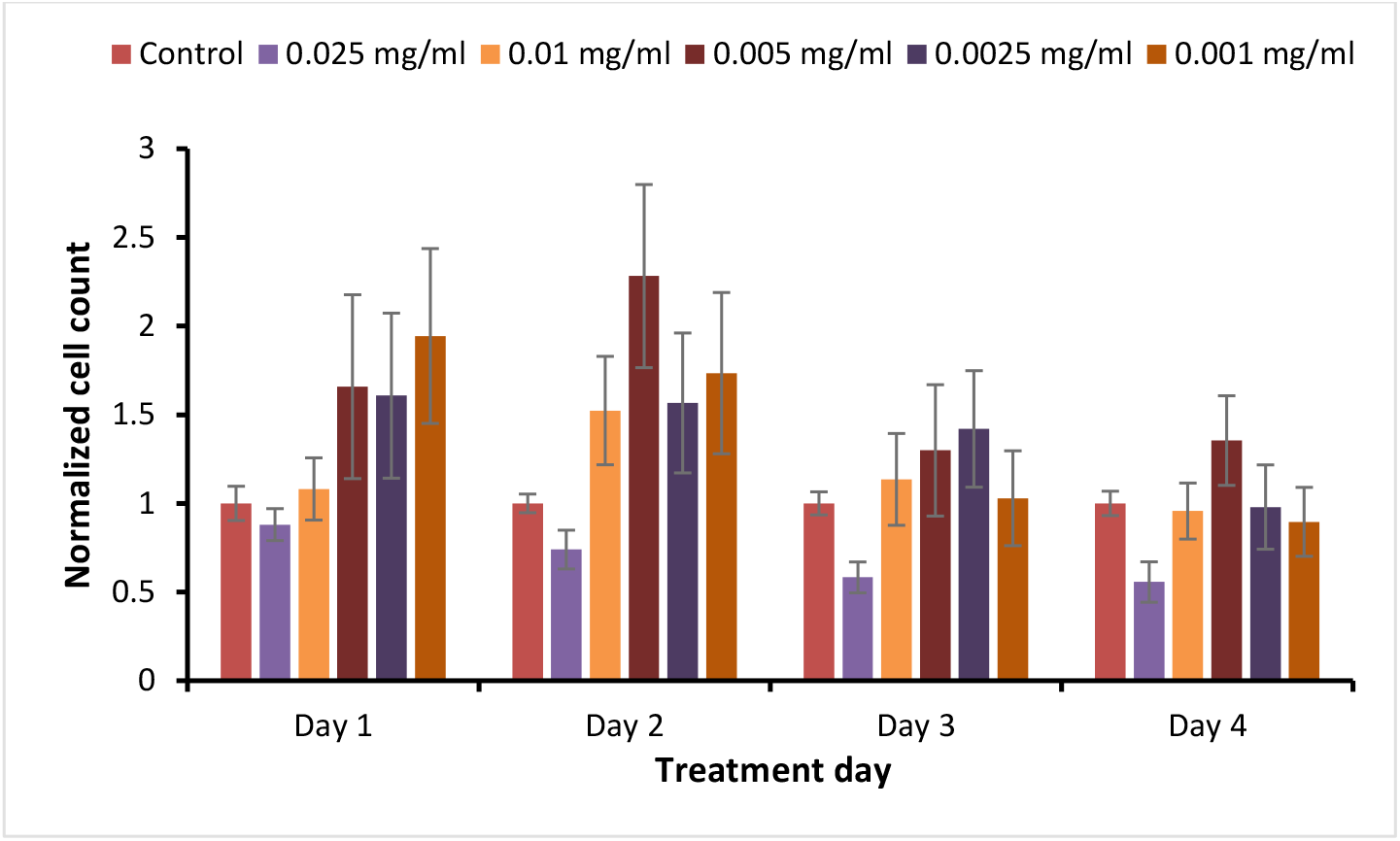
Proliferation assay for *P. africana* bark ethanol extract. Values are mean of three determinations (n = 3) ± SE.

**Figure 4:**
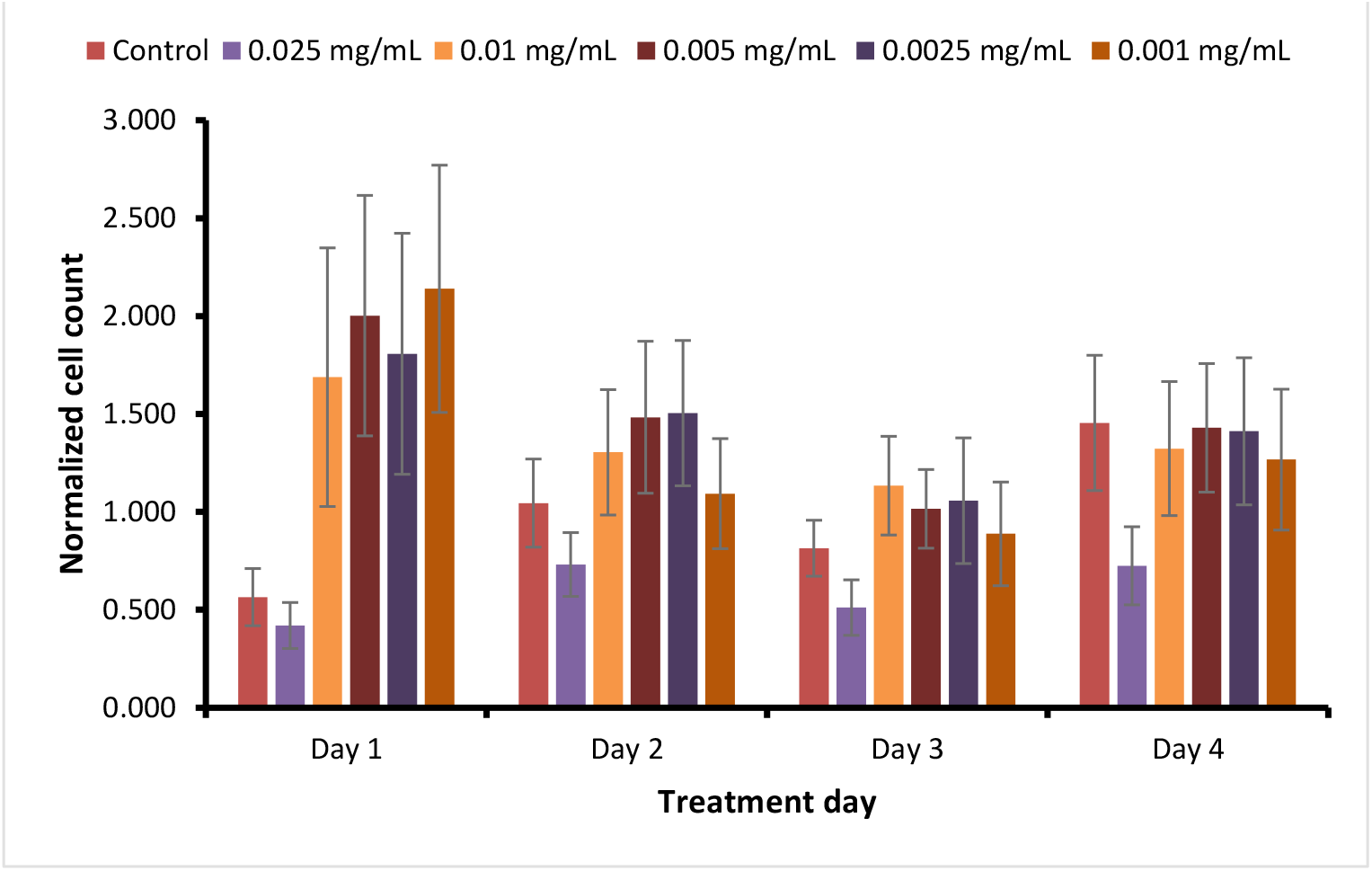
Proliferation assay for *P. africana* root methanol extract. Values are mean of three determinations (n = 3) ± SE.

Methanol is a more polar solvent than ethanol and this may be responsible for the difference in cell growth inhibition between the two extracts. This difference may be due to the fact that phenolics (suspected to play a major role in the cytotoxic effect of *P. africana* extracts) are often extracted in higher amounts in more polar solvents (Peschel et al., 2006; Qasim, Aziz, Rasheed, Gul, & Ajmal Khan, 2016; Sultana, Anwar, & Ashraf, 2009). This explanation is supported by the TPC and EC_50_ results of *P. africana* bark which were higher for methanol compared to ethanol extracts (Table 1, 2). Unlike methanol bark extracts, ethanol bark extracts did not show a significant difference between the control and the highest concentration tested, indicating that higher concentrations of ethanol extracts may be needed to achieve the same effect as the methanol extract (Figure 3). However, a clear dose-dependent growth inhibition of C4-2 cells treated with bark ethanol extracts can be seen.

**Table 2:**
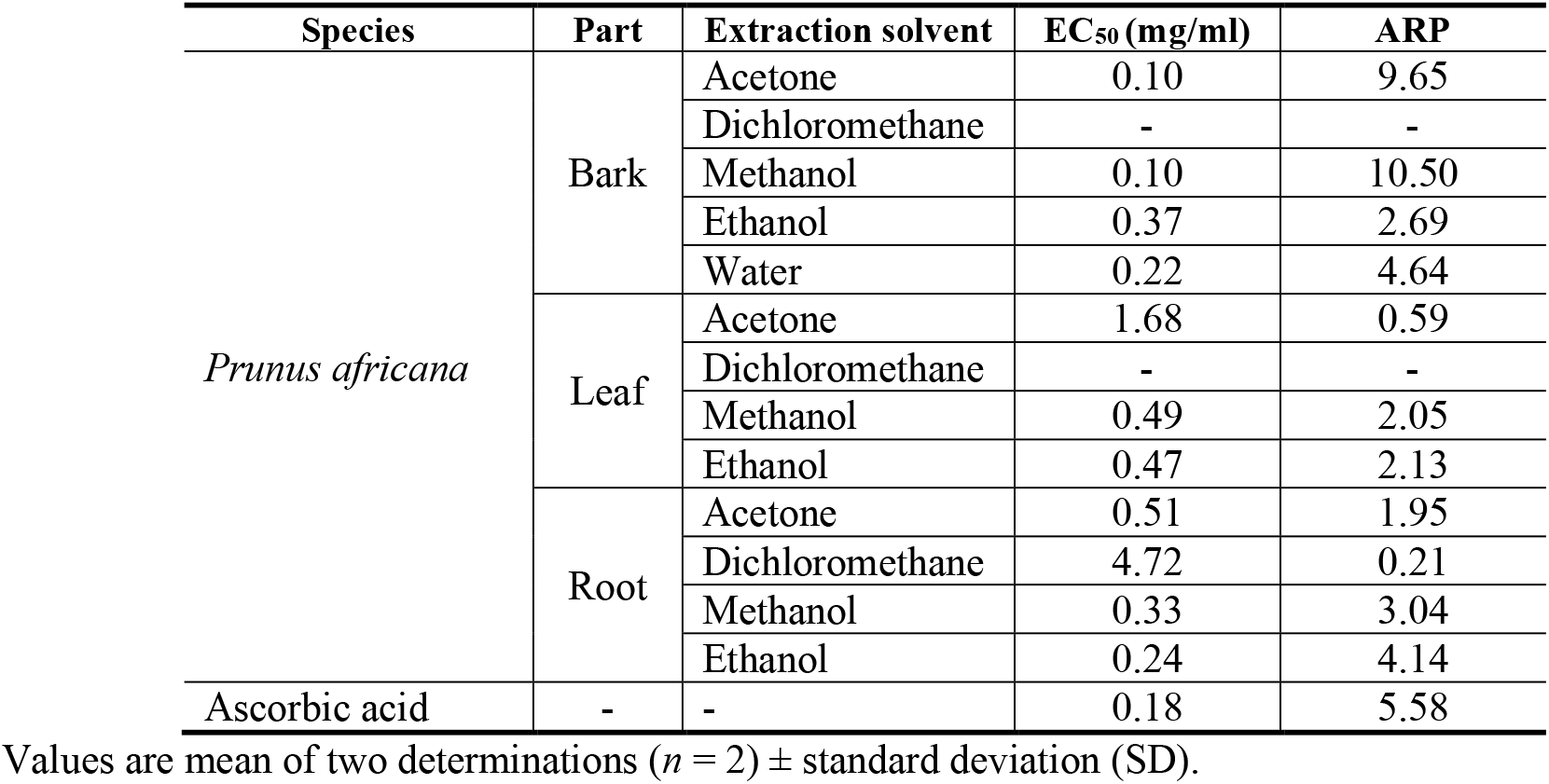
EC_50_ and Antiradical Power (ARP)

The growth inhibition effect of *P. africana* root extract is notable because although there have been a number of studies on the cytotoxic effects of *P. africana* bark extract, or components of bark extract, there have been few on the effects of other parts of the *P. africana* tree (Boulbès et al., 2006; Kadu et al., 2012; Karan et al., 2017; Komakech et al., 2017; Nabende, Karanja, Mwatha, & Wachira, 2015; Shenouda et al., 2007). Comparing extracts from the bark and root obtained by using the same solvent excludes any possible effect of extraction solvent on the extract composition and suggests that any differences seen are due to the natural composition of the plant parts. Although bark extracts had a higher inhibitory effect, the inhibitory effect of the root extracts was also evident, signifying that plant parts other than the bark can be utilized.

This is particularly important because *P. africana* bark has been exploited for use in the management of benign prostatic hypertrophy (BPH). Consequently, it is now an endangered species in Africa, needing to be conserved. This also has been recommended at international level by various organizations including UNESCO (Mugaka et al., 2013; Nabende et al., 2015). Utilization of other plant parts will hopefully lead to a less destructive exploitation of the *P. africana* tree, allowing for its use for medicinal purposes without the danger of extinction of the species. In addition, further analysis of other plant parts may lead to the discovery of additional secondary metabolites that may be useful for many other purposes. As more evidence emerges of the role secondary metabolites play in human health and nutrition, there is a need for more research into the composition of such medicinal plants and their potential uses. In light of the challenges facing chemotherapy and modern medicine such as drug toxicity and resistance, it has become imperative to find alternative drugs with improved specificity and efficiency.

## Conclusions

The first experimental study testing use of bark and root extracts of *P. africana* on C4-2 cells showed a concentration-dependent cytotoxicity, with the bark methanol extract showing the strongest effect. There was complete lysis of all cells above 0.05 mg/mL for all cells treated with bark and root extracts, while cells treated with leaf extracts showed a proliferation effect and clustering of cells compared to the control. The proliferation effect may be explained by the phenomenon hormesis, which is a dose-response phenomenon characterized by a low-dose stimulation and a high-dose inhibition or by a different composition of active secondary metabolites in the leaves than seen in the bark and root extracts. A four-day proliferation assay, using *P. africana* bark and root extracts at concentrations between 0.025 mg/mL and 0.001 mg/mL, showed growth inhibition of C4-2 cells in a dose dependent manner on all four days of the assay. There was a significant difference between control cells and cells treated with 0.025 mg/mL of the bark methanol extracts but not the bark ethanol or root methanol extracts. As C4-2 cells are hormonally insensitive and designed to mimic advanced prostate cancer, crude extracts of *P. africana* are a possible treatment option, not only for hormone sensitive prostate cancer, but also advanced, hormonally insensitive prostate cancer. Further research is however needed, to understand the mechanisms through which the extract causes inhibition of hormonally insensitive cancer.

## Acknowledgements

The authors gratefully acknowledge funding from USDA Evans-Allen which made this study possible.

## List of Abbreviations

AI: Androgen independent
AS: Androgen sensitive
BPH: Benign prostatic hyperplasia
DPPH: 2,2-diphenyl-1-picrylhydrazyl
DMSO: Dimethyl sulfoxide
EC_50_: 50% effective concentration
PBS: Phosphate buffered saline
RPMI: Roswell Park Memorial Institute media

